# Variability and reliability of effective connectivity within the core default mode network: A longitudinal spectral DCM study

**DOI:** 10.1101/273565

**Authors:** Hannes Almgren, Frederik Van de Steen, Simone Kühn, Adeel Razi, Karl Friston, Daniele Marinazzo

**Affiliations:** Department of Data Analysis, Faculty of Psychology and Educational Sciences, Ghent University, Belgium; Center for Lifespan Psychology, Max Planck Institute for Human Development, Berlin, Germany; Clinic and polyclinic for Psychiatry and Psychotherapy, University Clinic Hamburg-Eppendorf, Germany; The Wellcome Trust Centre for Neuroimaging, University College London, Queen Square, London WC1N 3BG, UK; Monash Institute of Cognitive and Clinical Neurosciences and Monash Biomedical Imaging, Monash University, Clayton, Australia; Department of Electronic Engineering, NED University of Engineering and Technology, Karachi, Pakistan

**Keywords:** Dynamic causal modelling, Resting State, fMRI, Effective connectivity, Reliability, Variability, Longitudinal designs

## Abstract

Dynamic causal modelling (DCM) for resting state fMRI – namely spectral DCM – is a recently developed and widely adopted method for inferring effective connectivity in intrinsic brain networks. Most research applying spectral DCM has focused on group-averaged connectivity within large-scale intrinsic brain networks; however, the consistency of subject- and session-specific estimates of effective connectivity has not been evaluated. Establishing reliability (within subjects) is crucial for its clinical use; e.g., as a neurophysiological phenotype of disease progression. Effective connectivity during rest is likely to vary due to changes in cognitive, behavioural, and physical states. Determining the sources of fluctuations in effective connectivity may yield greater understanding of brain processes and inform clinical applications about potential confounds. In the present study, we investigated the consistency of effective connectivity within and between subjects, as well as potential sources of variability (e.g., hemispheric asymmetry). We further investigated how standard procedures for data processing and signal extraction affect this consistency. DCM analyses were applied to four longitudinal resting state fMRI datasets. Our sample consisted of 20 subjects with 653 resting state fMRI sessions in total. These data allowed to quantify the robustness of connectivity estimates for each subject, and to draw conclusions beyond specific data features. We found that subjects contributing to all datasets showed systematic and reliable patterns of hemispheric asymmetry. When asymmetry was taken into account, subjects showed very similar connectivity patterns. We also found that various processing procedures (e.g. global signal regression and ROI size) had little effect on inference and reliability of connectivity for the majority of subjects. Bayesian model reduction increased reliability (within-subjects) and stability (between-subjects) of connectivity patterns.

## Introduction

During quiet wakefulness the brain shows several patterns of coherent activity, referred to as resting state networks (RSNs; Damoiseaux et al., 2006). RSNs include regions that are both functionally and structurally related (Van Den Heuvel, Mandl, Kahn, & Pol, 2009). Most studies investigating resting state networks are based on functional connectivity, which is defined as the statistical dependency among brain signals. However, interactions between brain regions are directed and are therefore not fully captured by (undirected) functional connectivity (Friston, 2011; Razi & Friston, 2016). Various methods have been developed to infer directed influences among brain regions, among which a prominent framework is Dynamic Causal Modelling (DCM; Friston, Harrison, & Penny, 2003).

DCM uses Bayesian model inversion procedures to estimate effective connectivity among neural populations from observed signals (e.g., BOLD-signals). It incorporates a biophysically plausible hemodynamic model (i.e., the Balloon model; Buxton, Wong, & Frank, 1998) to generate predicted BOLD-responses from neuronal states. Initially DCM was developed to estimate effective connectivity for experimental (task) fMRI studies (Friston et al., 2003). Recently, a DCM - referred to as spectral DCM (spDCM) - has been developed specifically to infer effective connectivity in resting state fMRI (Friston, Kahan, Biswal, & Razi, 2014). This DCM is based on a generative model of (complex) cross spectra between regional BOLD signals, and uses a power-law function (in the spectral domain) to model (random and endogenous) neuronal fluctuations. Fitting spectral (second-order) data features makes spDCM deterministic, which renders the estimation scheme computationally and statistically more efficient; compared to its stochastic counterpart that fits the (first-order) timeseries *per se* (i.e., stochastic DCM; Li et al., 2011).

The construct validity of spectral DCM has been established using both simulated and empirical data (Friston et al., 2014; Razi et al., 2015). Friston et al. (2014) simulated resting state fMRI timeseries for a network with three regions. Their results showed that spDCM estimates extrinsic effective connectivity with high accuracy, but tends to underestimate intrinsic connectivity (i.e., the inhibitory influence regions exert on themselves). In a subsequent *in silico* validation study, (Razi et al. 2015) demonstrated a similar accuracy for a network consisting of four regions. Interestingly, both studies showed that the root mean squared error (between the true and estimated connectivity) decreases with the number of scans. Both studies also showed that spDCM is sensitive for detecting group differences in effective connectivity. Most research using empirical data has focused on effective connectivity within the default mode network (e.g., Razi et al., 2015; Sharaev et al., 2016; Ushakov et al., 2016; Zhou et al. 2018). Both Razi et al. (2015) and Sharaev et al., 2016 estimated connectivity within the ‘core’ DMN, which included left and right intraparietal cortices (IPC), medial prefrontal cortex (mPFC), and posterior cingulate cortex (PCC). Both studies found reciprocal positive connectivity between IPC, and positive projections from lateral to medial brain regions. Ushakov et al. (2016) showed that adding extra regions (i.e., left and right parahippocampal gyri) to the four-region DMN did not have a substantial impact on its effective connectivity pattern. Zhou et al. (2018) showed that the salience and dorsal attention network have a negative influence on the core DMN, while the converse influence was slightly positive. Moreover, within the core DMN the same pattern of connectivity was found as in other spectral DCM studies.

In summary, studies that have applied spectral DCM to the DMN have yielded quite consistent results. However, these studies generally focused on group-averaged connectivity. While group studies are very useful to establish predictive validity, a thorough examination of subject and session-specific differences in effective connectivity during rest is an outstanding challenge. Quantifying within-subject stability is especially important in the context of single-patient diagnostics and predictions (Stephan et al., 2017). For other DCMs (e.g., DCM for task fMRI) test-retest reliability has been assessed between a few sessions, and was found to be good to excellent (e.g., Frässle, Paulus, Krach, & Jansen, 2015; Schuyler et al., 2010).

Here, we wanted to assess within-subject reliability (and between-subject consistency) of effective connectivity estimated by *spectral* DCM across many resting state fMRI sessions acquired in longitudinal studies. Although effective connectivity during resting state fMRI should be sufficiently reliable to be used in a clinical context, it is likely to vary as a consequence of changes in physical, emotional and behavioural states (e.g., amount of sleep), and the sources of this variability need to be established. Assessment of these longitudinal variations in effective connectivity could yield important insights in the effects of behavioural and psychological states on macroscopic brain dynamics (see, e.g., Laumann et al., 2015). The goals of the present study were to assess whether, and to what extent, connectivity patterns in the default mode network are consistent both within and between subjects, and to investigate the sources of variability in effective connectivity. To meet these aims, we made use of four longitudinal datasets, with a minimum of ten resting state sessions for each subject. These datasets allowed us to quantify the stability of the posterior estimates of (effective) connectivity in the default mode network across sessions, and to generalize conclusions beyond specific datasets (e.g., subjects’ characteristics, scanning parameters, *etc*).

## Methods

### Datasets and subjects

Data were obtained from four extensive longitudinal datasets acquired at different research institutions. The total sample consisted of 20 subjects (11 females, age range at start of studies: 24 – 45 years) with a minimum of 10 resting state sessions for each subject. Altogether, the datasets contained 653 rsfMRI sessions. A summary of the datasets is shown in Table 1.

**Table 1:** *subject information.* ^1^Age at the start of study,^2^Total number of sessions initially included in the present study (these differ slightly from the original studies because some low-quality images were not shared, duplicates were encountered, or initial pilot sessions were not included), ^3^Span = approximate time over which all rsfMRI scan sessions took place. ^4^Total number of scans in each session. Abbreviations: MyConn = Myconnectome, MSC = Midnight Scan Club.

| Subject | Dataset | M/F | Age<sup>1</sup> | Total sessions<sup>2</sup> | Span<sup>3</sup> | Total scans (time)<sup>4</sup> |
| --- | --- | --- | --- | --- | --- | --- |
| 1 | MyConn | M | 45 | 89 | ±1.5 years | 518 (±10 min) |
| 2 | Kirby | M | 40 | 156 | ±3.5 years | 200 (±7 min) |
| 3 | Day2day | F | 24 | 50 | ±5.5 months | 150 (±5 min) |
| 4 | Day2day | F | 28 | 13 | ±3.5 months | 150 (±5 min) |
| 5 | Day2day | F | 31 | 50 | ±13 months | 150 (±5 min) |
| 6 | Day2day | M | 32 | 11 | ±2 months | 150 (±5 min) |
| 7 | Day2day | F | 29 | 45 | ±7 months | 150 (±5 min) |
| 8 | Day2day | F | 24 | 47 | ±5.5 months | 150 (±5 min) |
| 9 | Day2day | M | 30 | 43 | ±7 months | 150 (±5 min) |
| 10 | Day2day | F | 29 | 49 | ±7.5 months | 150 (±5 min) |
| 11 | MSC | M | 34 | 10 | 17 days | 818 (±30 min) |
| 12 | MSC | M | 34 | 10 | 10 days | 818 (±30 min) |
| 13 | MSC | F | 29 | 10 | 12 days | 818 (±30 min) |
| 14 | MSC | F | 28 | 10 | 15 days | 818 (±30 min) |
| 15 | MSC | M | 27 | 10 | 14 days | 818 (±30 min) |
| 16 | MSC | F | 24 | 10 | 15 days | 818 (±30 min) |
| 17 | MSC | F | 31 | 10 | 39 days | 818 ($\pm 30$ min) |
| 18 | MSC | F | 27 | 10 | 18 days | 818 ($\pm 30$ min) |
| 19 | MSC | M | 26 | 10 | 16 days | 818 ($\pm 30$ min) |
| 20 | MSC | M | 31 | 10 | 21 days | 818 ($\pm 30$ min) |

*Dataset 1.* The first dataset (‘myconnectome’) was part of the MyConnectome project (see, Laumann et al., 2015). During this project, resting state fMRI scans were acquired from a single person (male, 45 years at the start of study) on 89 occasions over the period of 1.5 years. MRI data were obtained with a Siemens MAGNETOM Skyra 3T MRI scanner (Siemens, Erlangen, Germany), using a 32-channel head coil. Resting state fMRI was acquired using a multi-band echo-planar imaging (MBEPI) sequence (TR = 1160ms; TE = 30ms; voxel size = 2.4mm x 2.4mm x 2mm; FOV = 230mm; flip angle = 63 degrees; multi-band factor = 4). Functional images for the first 14 sessions contained 68 slices; images from remaining sessions comprised 64 slices. Resting state scan length was approximately 10 minutes (518 images). T1 images were acquired using a MPRAGE sequence (TR = 2400ms; TE = 2.14ms; TI = 1000ms; voxel size = 0.8mm isotropic; 256 sagittal slices; flip angle = 8 degrees; GRAPPA factor = 2). Only the T1-weighted image acquired during the session prior to the first rsfMRI session was used in the present study (i.e., for coregistration and normalization). This dataset was obtained from the OpenfMRI database. Its accession number is ds000031.

*Dataset 2.* The second dataset (‘Kirby’) contained data acquired from a single subject (40 years at the start of study, male) on 156 occasions (3.5 years; see, Choe et al., 2015). The subject was scanned using a 3T Philips Achieva scanner (Philips Healthcare, Best, Netherlands), with a 16-channel neurovascular coil. Functional resting state data were acquired using a multi-slice SENSE-EPI sequence (TR = 2000ms; TE = 30ms; voxel size = 3mm x 3mm x 3mm; flip angle = 75 degrees; 37 axial slices; SENSE factor = 2). Scan length was approximately 7 minutes (200 images). T1-weighted images were acquired using a MPRAGE sequence (TR = 6.7ms; TE = 3.1ms; TI = 842ms; voxel size = 1.0mm x 1.0mm x 1.2mm; flip angle = 8 degrees; SENSE factor = 2). The T1-weighted image acquired during the first scan session was used in the present study.

*Dataset 3.* The third dataset (‘day2day’) contained data acquired from eight subjects (6 females; age range 24 - 32 years; Filevich et al., 2017). The number of scan sessions per subject ranged from 11 to 50 (sessions in total), and were acquired within a period of 2 to 13 months. Subjects were scanned using a 3T Magnetom Trio MRI scanner (Siemens, Erlangen, Germany) and a 12-channel head coil. RsfMRI data was acquired using a T2*-weighted echo planar imaging (EPI) sequence (TR = 2000ms; TE = 30ms; voxel size = 3mm x 3mm x 3mm; flip angle = 80 degrees; 36 axial slices; GRAPPA acceleration factor = 2). The length of resting state scanning was approximately five minutes (150 images). Structural MRI scans were acquired using a MPRAGE sequence (TR = 2500ms; TE = 4.77ms; TI = 1100ms; voxel size = 1.0mm x 1.0mm x 1.0m; flip angle = 7 degrees). Only the T1-weighted image acquired during the first scan session was used.

*Dataset 4.* The fourth dataset (‘midnight scan club’) contained data from ten subjects (5 females; age range 24 – 35 years; see, Gordon et al., 2017). Participants were scanned at midnight on twelve consecutive days with a 3T Siemens Trio MRI scanner (Siemens, Erlangen, Germany). On ten occasions, rsfMRI data were acquired with a gradient-echo EPI sequence (TR= 2200ms; TE = 27ms; voxel size = 4mm x 4mm x 4mm; flip angle = 90 degrees; 36 axial slices). Each session contained 818 volumes (approximately 30 minutes). Structural scans were acquired using a gradient-recalled inverse recovery (GR-IR) sequence (TR = 2400ms; TE = 3.74ms; TI = 1000ms; voxel size = 0.8mm x 0.8mm x 0.8mm; 224 sagittal slices; flip angle = 8 degrees). The first structural image acquired from each subject was used for the analyses. This data was obtained from the OpenfMRI database. Its accession number is ds000224.

## Data Analyses

### Preprocessing

Preprocessing was performed using the SPM12 software package (revision 6906; Wellcome Centre for Human Neuroimaging; www.fil.ion.ucl.ac.uk/spm/software/spm12). The first five images of each session‘s rsfMRI sequence were discarded to allow for T1 equilibration. First, resting state fMRI images were corrected for differences in slice timing (using the central slice of each volume as a reference). Next, images were realigned to the first functional volume of each session. Images were then coregistered to the skull-stripped anatomical image. Finally, images were normalized to MNI space (Montreal Neurological Institute) and smoothed using a Gaussian kernel (6mm FWHM).

### Time-series extraction

Session-specific DMN voxels were identified by specifying and estimating a GLM containing: (1) a discrete cosine basis set as principal regressor (frequency range: 0.0078 – 0.1Hz), (2) six head motion regressors (three translational, three rotational), (3) a regressor for CSF signal (principal eigenvariate of 5mm ROI within CSF circulation system), and (4) a regressor containing WM signal (principal eigenvariate of 7mm ROI within brainstem). The number of cosine components was related to the number of scans within a session. An F-contrast was specified across all DCT components to produce an SPM, which was masked using ROIs (sphere radius = 10mm) extracted from template ICA maps (Smith et al., 2009). ROI centre coordinates were (x=2; y=-58; z=30) for precuneus, (x=2; y=56; z=-4) for medial prefrontal cortex, (x=-44; y=-60; z=24) for the left inferior parietal cortex, and (x=54; y=- 62; z=28) for the right inferior parietal cortex (see, Figure 1: left panel). Coordinates were labelled using the AAL atlas. Time-series were acquired by computing the principal eigenvariate of signals from voxels centered on the peak voxel of the aforementioned F-contrast (session-specific; sphere radius = 8mm) within each ROI, which allowed session- and subject-specific differences in exact location of DMN regions. Voxels were only included if they survived an a priori specified threshold: if sessions contained less than 200 scans per session, voxels were included if they exceeded an uncorrected (full brain) alpha-threshold of 0.05. If sessions contained more scans, a stepwise increase in alpha-threshold (alpha = 0.001, 0.01, to 0.05) was applied, until significant voxels were detected (using an upper boundary of alpha = 0.05). Importantly, the alpha-level specified here is used to detect voxels that contain low-frequency fluctuations, and is independent from the criterion used to infer connectivity.

**Figure 1.**
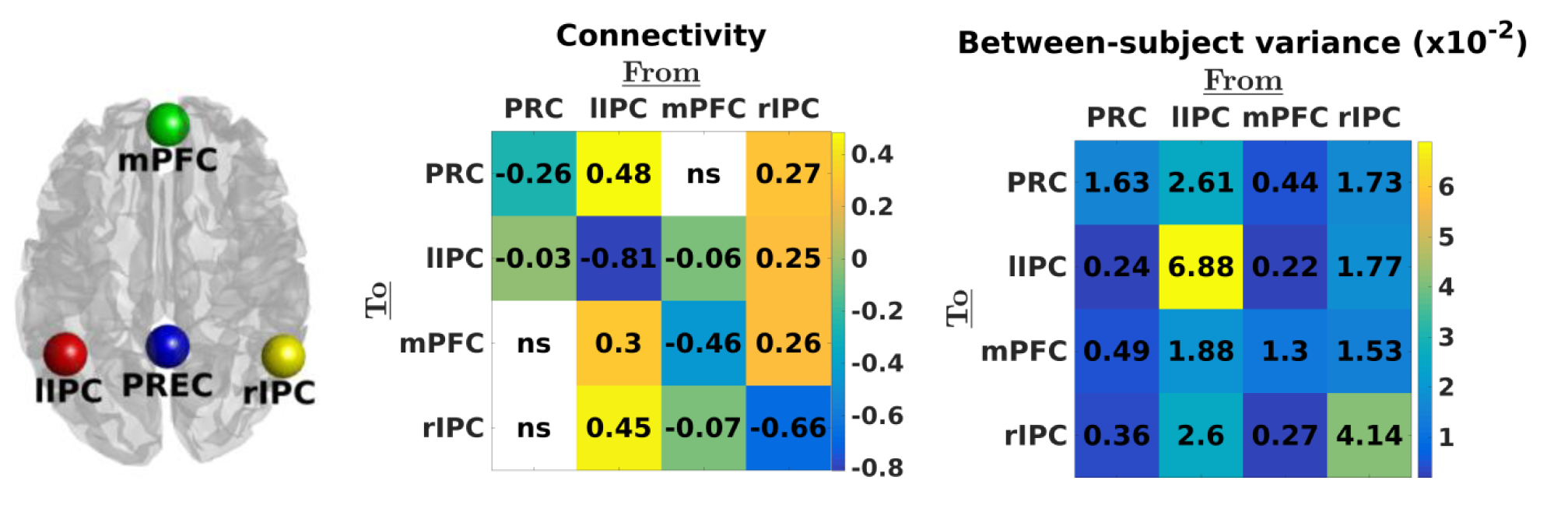
Left panel: location of ROIs used in the present study. Middle panel: Estimated effective connectivity (from columns to rows) at the group level. Diagonal elements reflect self-inhibition parameterised in log-scale (relative to the prior mean of −0.5Hz). A posterior probability criterion of 90% was used. Right panel: estimated between-subject variability for each connection (PEB.Ce). It is evident that the left and right IPC showed the greatest between-subject variability in self-inhibition (log-scale) and extrinsic connectivity (hertz).

### (Spectral) Dynamic Causal Modelling and Hierarchical Bayes

Fully-connected DCMs were specified and inverted for each session separately (DCM12; revision 6801), without the specification of exogenous (i.e., experimental) inputs. Sessions were excluded from all analyses if they did not meet five diagnostic criteria, namely (1) explained variance of predicted BOLD signals above 60%, (2) at least one connection with a connection strength greater than 1/8 Hz, (3) at least one effectively estimated parameter (based on Kullback-Leibler divergence of posterior from prior distribution), (4) maximum frame-wise displacement (FD) under 1.5mm, and (5) a maximum alpha-threshold of 0.05 for which significant voxels were found in all DMN regions for that session. Three subjects (i.e., S13, S18, and S19) were rejected because they had less than 8 sessions after rejection based on diagnostic checks. Additionally, thirty-one sessions (across subjects and datasets) were excluded because they did not meet diagnostic criteria, and three sessions were discarded because of other problems (e.g., incomplete volumes). A total of 589 rsfMRI sessions were included after quality and diagnostic checks.

To compute average connectivity at the subject-level we specified Parametric Empirical Bayes (PEB; Friston et al., 2016) models with one regressor (comprising a column of ones) that modelled average (within-subject) connectivity over sessions. The subject-specific PEB models were then included in a group-level PEB model, including again one regressor to compute average (between-subject) connectivity over subjects. Default settings were used for estimation at the subject and group level; i.e., the prior covariance of connections had the same form at the session, subject and group-level, where the within subject (between session) prior covariance was 1/16^th^ of the prior covariance of (between-subject) group means. However, every parameter was equipped with a separate between-subject precision component. Only connectivity-related parameters (‘A-matrix’) were included as dependent variables in subject and group analyses. All inference was made using a posterior probability criterion of 90% for each connection.

### Stability criteria

Stability of the strength and direction of connections was assessed for three different network characteristics. (1) We tested hemispheric asymmetry for each session and subject by computing the posterior probability that the average outgoing connectivity (i.e., effluent or out-degree) from the left IPC differed from the average outgoing connectivity from the right IPC (posterior probability criterion = 90%). Between (resp., within) subject stability of asymmetry was evaluated by computing the ratio of subjects (resp., sessions) that showed most prominent influence from either right or left IPC. (2) We assessed stability of the estimated connectivity matrices (i.e., over all connections) by calculating the average correlation between vectorised connectivity matrices for each pair of subjects and sessions. (3) We assessed between (resp., within) subject stability of the type of connection (i.e., excitatory, inhibitory, or no influence) by computing the percentage of subjects (resp., sessions) that showed either a positive, negative, or non-existent influence between regions (using the 90% posterior confidence criterion). The latter stability measure was computed for each connection separately.

### Effect of (pre)processing steps

The effect of three (pre)processing steps on connectivity and stability was assessed. (1) (Empirical) Bayesian model reduction was applied to assess the effect of using subject-specific and group connectivity as empirical priors for session- and subject-specific estimation, respectively. We compared stability with and without empirically optimising connectivity parameters at the session and subject level (as implemented by spm_dcm_peb.m). (2) To assess the effect of ROI size on connectivity and stability, we reanalyzed the data using spherical ROIs with radii of 4mm, 8mm, 12mm and 16mm. ROIs were cantered at the average coordinate of the (session-specific) voxels included in the previous ‘basic’ analyses. To allow proper comparison, we calculated the eigenvariate of all significant voxels in the sphere (i.e., without using the conjunction with the ROI derived from the ICA template). The same subjects were excluded as in the basic analyses. To ensure proper comparison between analyses with different ROI sizes, sessions were excluded if they did not reach diagnostic thresholds for all ROI size. Consequently, 57 sessions were discarded, which yielded a total sample of 563 sessions. (3) Finally, we assessed the influence of global signal regression (GSR) on the reliability and connectivity. Therefore we repeated the ‘basic’ analyses with GSR, which was done by scaling (preprocessed) fMRI volumes with the inverse of the scan-specific global mean intensity (in SPM: global normalization = ‘scaling’). All subsequent analyses (e.g., peak-value coordinate detection, time-series extraction) were performed using these scaled images. Again, same subjects were excluded as in the basic analyses. Forty-two sessions were excluded because they did not reach diagnostic thresholds for one or both analyses (i.e., with or without GSR), which left a sample of 578 rsfMRI sessions.

## Results

### Group-level

Estimated connectivity at the group level is shown in Figure 1. Clearly, connections from bilateral IPC were stronger compared to other connections. Moreover, the left IPC showed a stronger outgoing influence compared to right IPC (mean difference = 0.15; SD = 0.03; posterior probability > 0.99) and the lowest self-inhibition. Between-subject variability for both extrinsic (i.e., between-regions) and intrinsic connections (i.e., self-inhibition) was greater for projections arising from left and right IPC compared to other regions.

### Between-subject variability and consistency

Average connectivity for some exemplar subjects is shown in Figure 2. Most subjects showed a dominant influence from either left or right IPC. Post-hoc tests comparing average outgoing connectivity from left IPC to average connectivity from right IPC indeed confirmed our observations: ten subjects showed higher influence from left IPC, while six subjects showed greater influence from right IPC. The average difference between left and right IPC connectivity was 0.60Hz, indicating a non-trivial effect. The dominant IPC showed lowest self-inhibition in fifteen out of sixteen asymmetric subjects. To assess the similarity in general connectivity patterns between subjects, we calculated correlations between (vectorised) connectivity matrices for all possible pairs of subjects (136 pairs in total). We accommodated hemispheric asymmetry by swapping the columns and rows representing left and right IPC for all left-asymmetric subjects. This yielded reordered matrices for which the second column and row represented the dominant hemisphere and the fourth column and row represented the non-dominant IPC in all subjects. The results of this analysis are shown in Figure 3. The ensuing average correlation was 0.56 (range: [−0.31 0.96]) for the original and 0.81 (range: [0.45 0.98]) for the reordered connectivity matrices. In short, casting effective connectivity in terms of (out-degree) *dominant* versus non-*dominant*, as opposed to *right* versus *left*, hemisphere markedly improved between-subject consistency. The predominance of asymmetry in between-subject variability was also confirmed with a principal component analysis (PCA) on effective connectivity across subjects. The first principal component showed highest (and opposite) loadings on left and right IPC, and explained approximately 62% of total variance. All subsequent between-subject analyses were performed using dominance-ordered connectivity matrices. To assess consistency in terms of connection type, we enumerated (for each connection separately) the number of subjects showing excitatory, inhibitory, or non-existent influence (i.e., with a posterior probability < 90%). The results are shown in Figure 4. The dominant IPC exerted a positive influence on all other regions for all subjects, while the influence of the non-dominant IPC varied between subjects. Moreover, influence from precuneus on mPFC was significantly positive in 11 out of 17 subjects (64.7%) and connectivity from the mPFC to dominant IPC was negative in 12 out of 17 subjects (70.6%).

**Figure 2.**
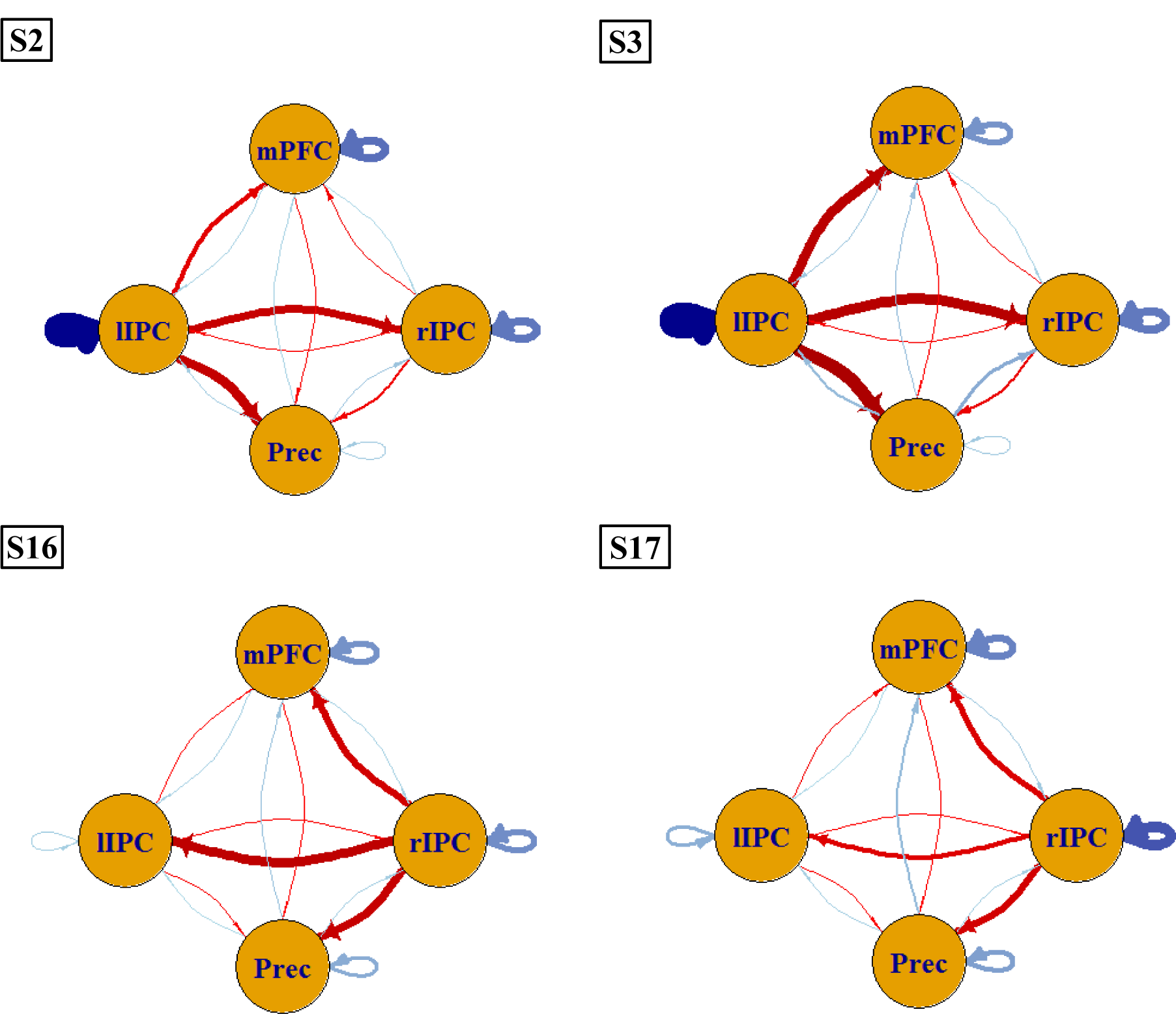
Connectivity patterns for exemplar subjects. For extrinsic connections between regions, red lines denote positive connectivity and blue lines negative connectivity. For self-connections, red lines depict connectivity above the prior mean, while blue lines depict connectivity lower than the prior mean (i.e., −0,5 Hz). Across datasets, subjects showed most dominant influence from either left (e.g., S2 and S3) or right IPC (e.g., S16 and S17). Moreover, self-inhibition was lowest for the dominant IPC. Line thickness and brightness reflect the strength of the respective connection.

**Figure 3.**
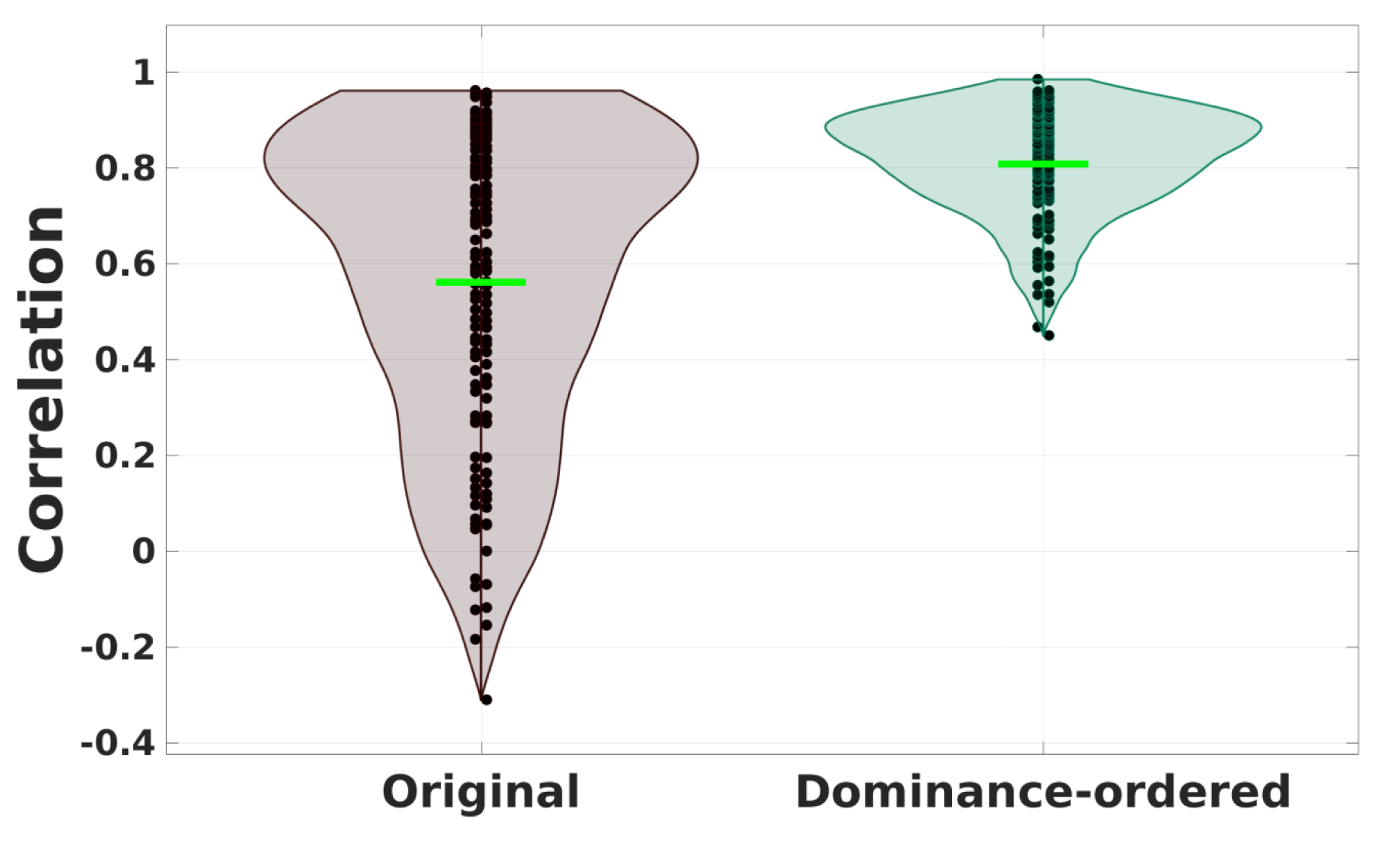
Violin plots of correlations between original (left plot) and dominance-ordered (right plot) connectivity-matrices for all possible pairs of subjects. Horizontal green lines depict the mean correlation. Clearly, the consistency is much higher when hemispheric dominance is taken into account.

**Figure 4.**
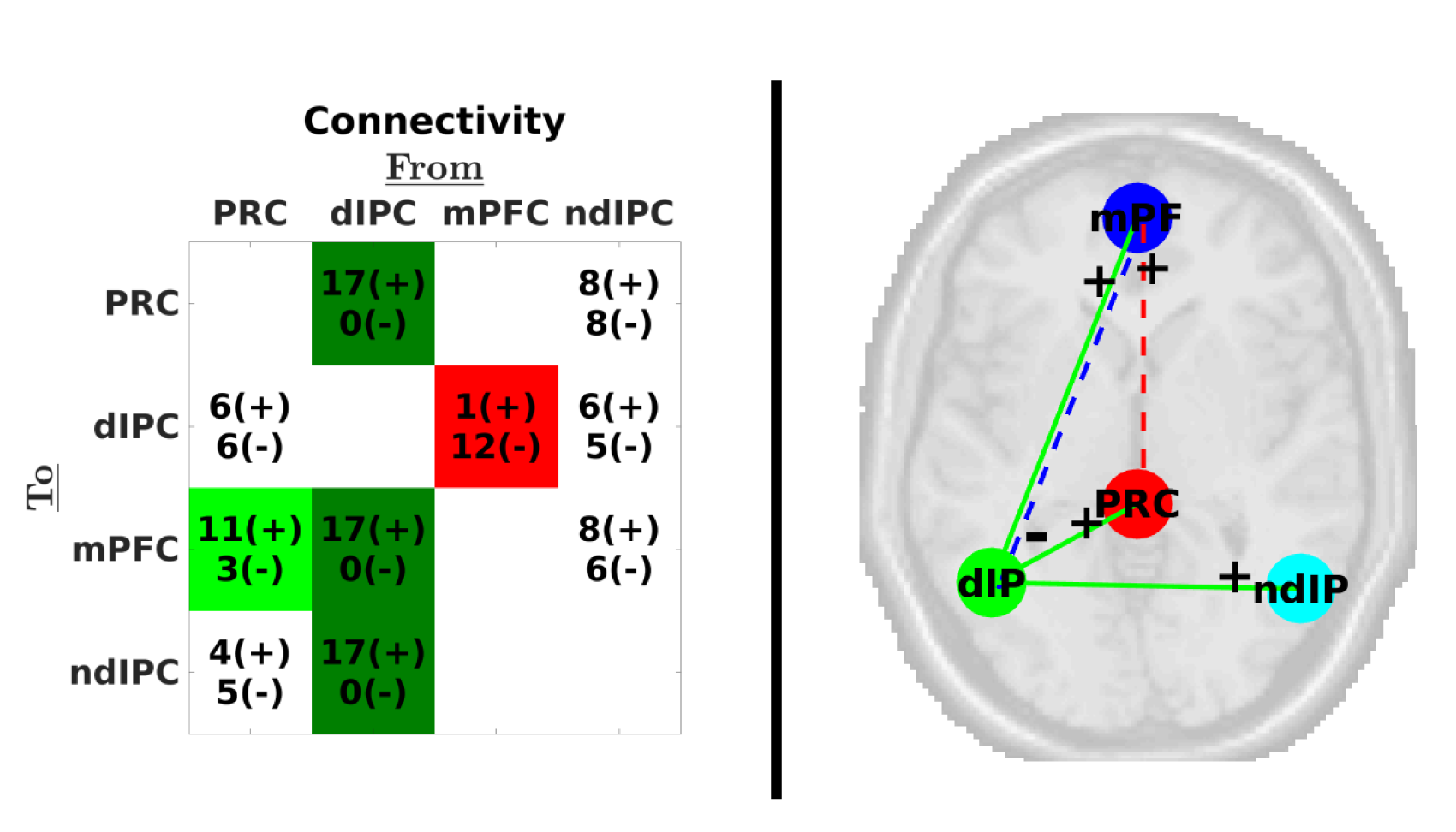
Left panel: Number of subjects with excitatory (positive) or inhibitory (negative) influences (posterior probability > 90%; effective connectivity from column to row regions). Connections showing positive influence for at least 75% (i.e., 13 out of 17) of subjects are shown in dark green, connections showing positive influence for at least 60% (i.e., 11 out of 17) of subjects are shown in light green. Connections showing negative influence for at least 60% of subjects are shown in red. Self-connections are omitted for simplicity. Right panel: Thresholded network showing consistent connectivity types (connections showing same influence for at least 60% of participants are shown). Full lines depict connections with the same sign in all subjects, dashed lines depict connections showing same sign in at least 60% of participants. For visualization purposes the precuneus is shown more anteriorly than in reality. Anatomical labels: PRC = precuneus; mPF = medial prefrontal cortex; dIP = dominant inferior parietal cortex; ndIP = non-dominant inferior parietal cortex.

### Within-subject variability and reliability

The same tests for hemispheric asymmetry (i.e., comparing average connectivity from left versus average connectivity from right IPC) were performed on every session‘s connectivity matrix. Results for all subjects are shown in Figure 5. On average, 72% (range: [0.4 0.93]) of sessions showed the same asymmetry as the respective subject-specific asymmetry. Most subjects showed reliable asymmetry, while some subjects showed a more variable asymmetry. PCA revealed that the first principal component of 8 subjects showed opposite loadings on left compared to right IPC, accounting on average for 40% (range: 24-73%) of between-session variance. Additionally, two subjects showed asymmetric loadings for the second or third principal component, accounting on average for 22% of variance (range: 20-23%). The reliability of general connectivity patterns was assessed by computing the average correlation across all possible pairs of sessions for both original and reordered matrices. Figure 6 shows the results of these analyses. The average correlation was 0.39 (range: 0.08 – 0.71) for the original matrices, and 0.51 (range: 0.13 – 0.79) for dominance-ordered matrices. In subsequent analyses we assessed the sign-stability of connections within subjects. Figure 7 shows connections that had the same influence (i.e., inhibitory or excitatory) in at least 75% of sessions for exemplar subjects. Subjects showed notably stable positive connectivity from either left or right IPC (e.g., subject 3 and subject 16, respectively), which nicely coincides with the average subject-specific hemispheric asymmetry reviewed in the previous paragraph. Additionally, each subject showed unique stable connections.

**Figure 5.**
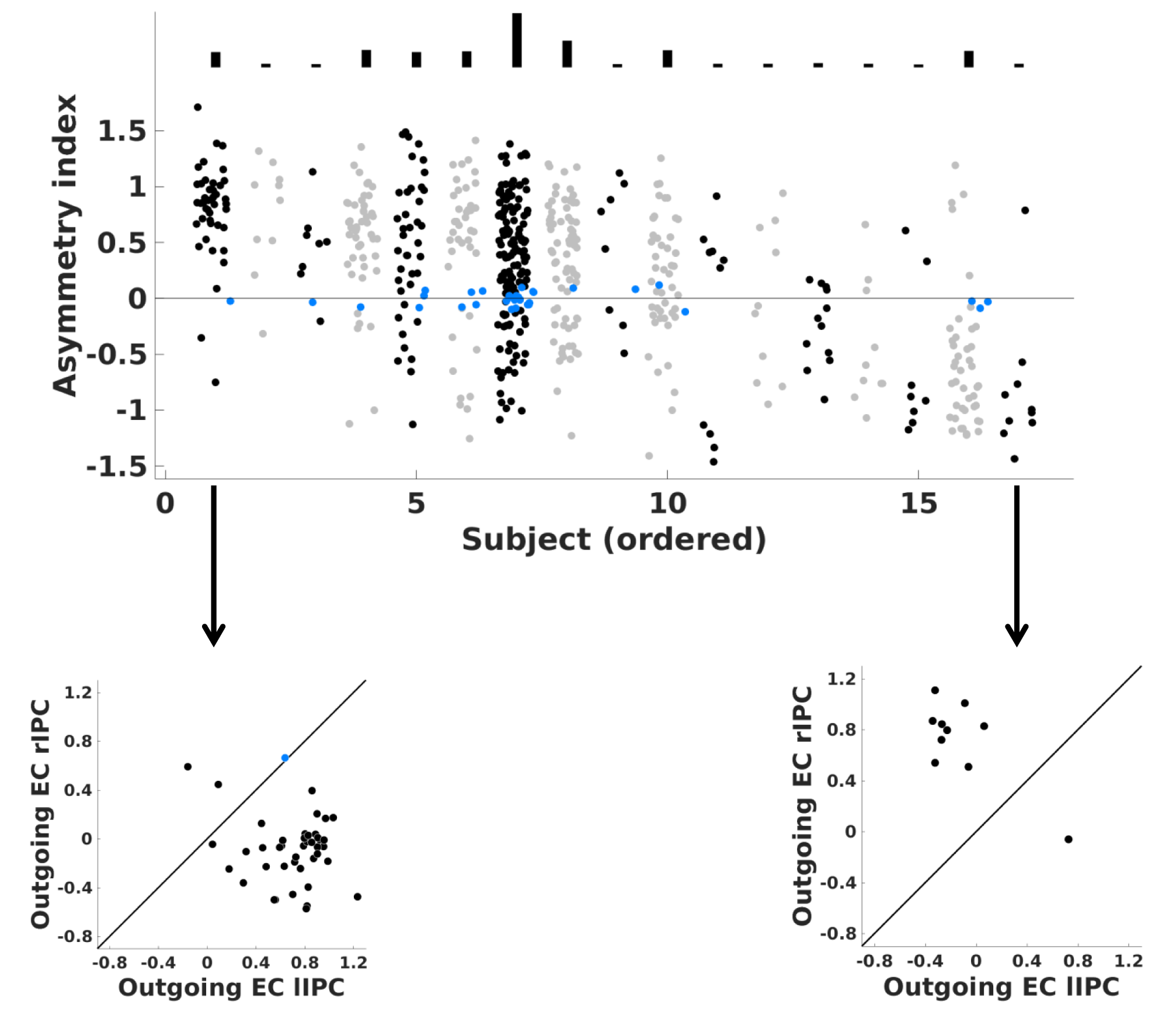
Upper panel: Session-specific hemispheric asymmetry for all subjects, in order of decreasing proportion of left hemisphere-dominant sessions. Dots represent the asymmetry index for each session of the respective subject (positive = left dominant; negative = right dominant); bars above the figure depict the number of sessions for each subject. Blue dots depict sessions without evidence for hemispheric dominance. Lower panels: Hemispheric asymmetry for the most stable left and right asymmetric subject. Black circles represent the average outgoing influence from left (x-axis) and right IPC (y-axis). Circles below the reference line indicate sessions with higher influence from left IPC, circles above the reference line depict sessions with higher influence from right IPC. Light-blue circles depict sessions for which asymmetry did not survive the posterior probability criterion.

**Figure 6.**
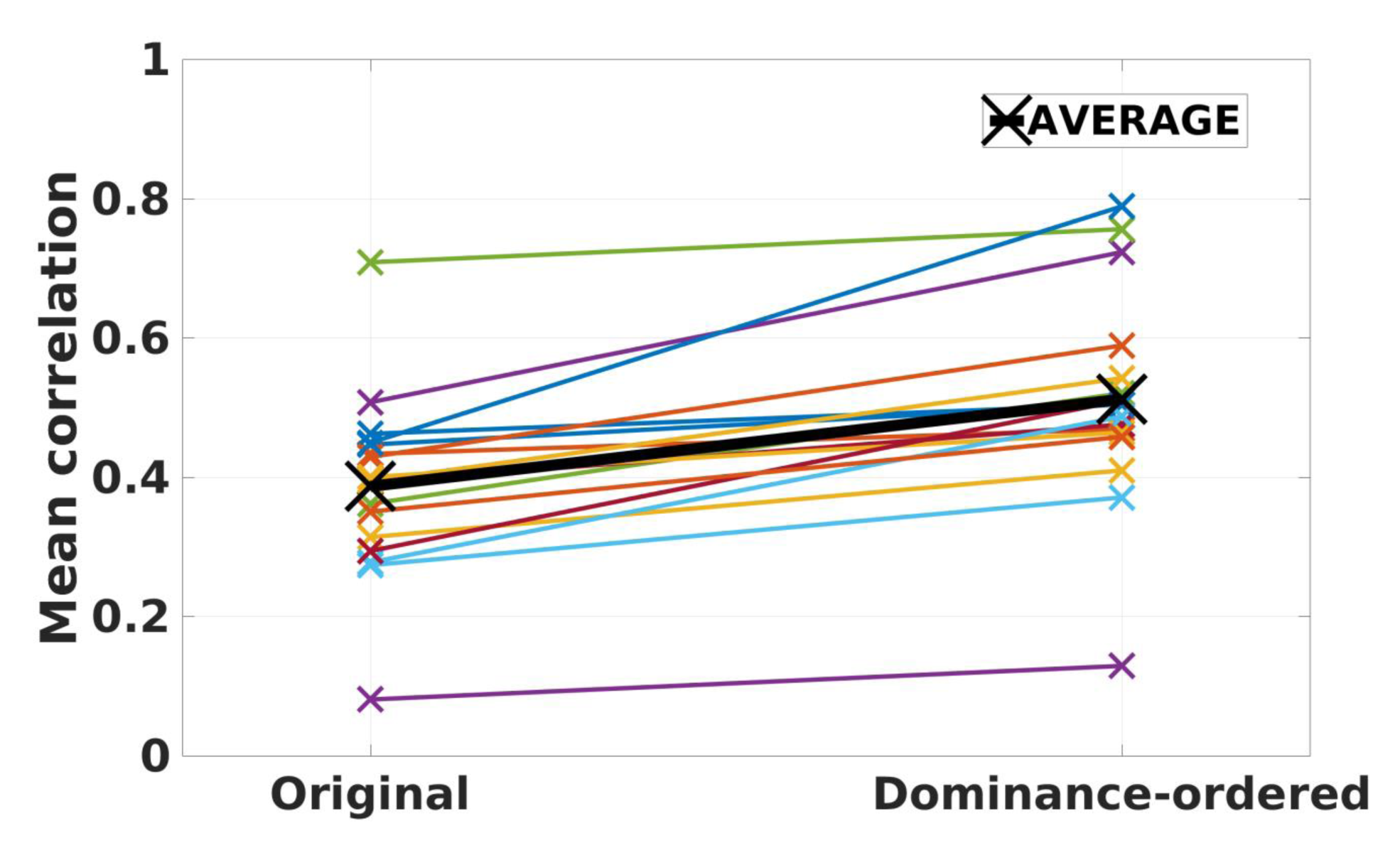
Average correlation across all possible pairs of sessions, for each subject (represented with colored lines) using original (left) and dominance-ordered (right) connectivity matrices. For each subject, the correlation increased slightly after accounting for hemispheric asymmetry. The bold black line represents the average correlation across subjects: 0.39 for the original and 0.51 for dominance ordered matrices.

**Figure 7.**
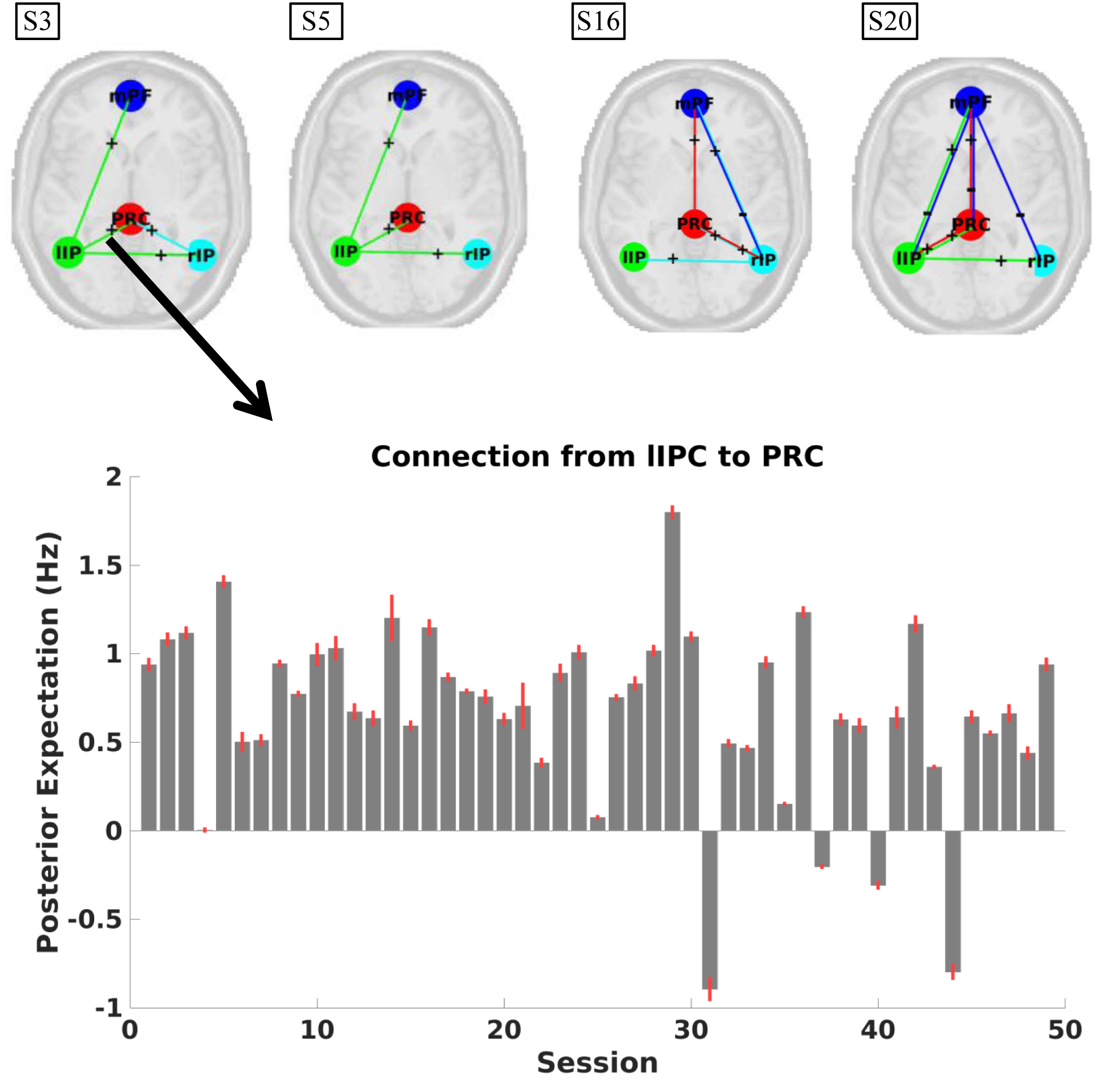
Upper panel: Connections having the same sign in at least 75% of sessions within the respective (exemplar) subject. The line colours depict the source of a connection (e.g., green lines depict connections from the lIPC to other regions). Clearly, stable connections arise from left or right IPC, which nicely coincides with the subject-specific asymmetry. For visualization purposes the precuneus is shown more anteriorly than in reality. Lower panel: Posterior estimates of the strongest connection for subject 3, plotted against session number.

### Effect of (Pre)processing

#### Bayesian Model Reduction (BMR)

At the subject level, hemispheric asymmetry was not changed (i.e., from left to right or vice versa) in any subject after including empirical priors. Consistency of connectivity patterns between subjects increased after empirical BMR for both original and reordered matrices (increase in average correlation after empirical BMR was 10.2% for original matrices and 5.6% for reordered matrices). The consistency of connectivity (within-subjects) increased for 15 out of 17 subjects using original connectivity matrices and for 13 out of 17 subjects using reordered connectivity matrices (average increase in correlation was 15.4% for original and 9.8% for reordered matrices).

#### Effect of ROI size

Next, we assessed the influence of ROI size on connectivity at the group- and subject-level, as well as its influence on reliability. Group-level results are shown in Figure 8. Generally, connectivity patterns were very similar for different ROI sizes at the group-level (mean correlation = 0.980; range = 0.954-0.997). Asymmetry decreased with increasing ROI size (see Figure 8, lower panel), but was left dominant, even for larger ROI sizes (posterior probability > 0.90). At the subject level (not shown), hemispheric asymmetry did not change for any ROI size in 11 subjects (65%), while in three subjects (18%) hemispheric asymmetry flipped for some ROI sizes. The consistency of connectivity, assessed as the average correlation between effective connectivity matrices, was 0.40, 0.39, 0.43, and 0.41 for increasing ROI radii (i.e., 4, 8, 12, and 16mm).

**Figure 8.**
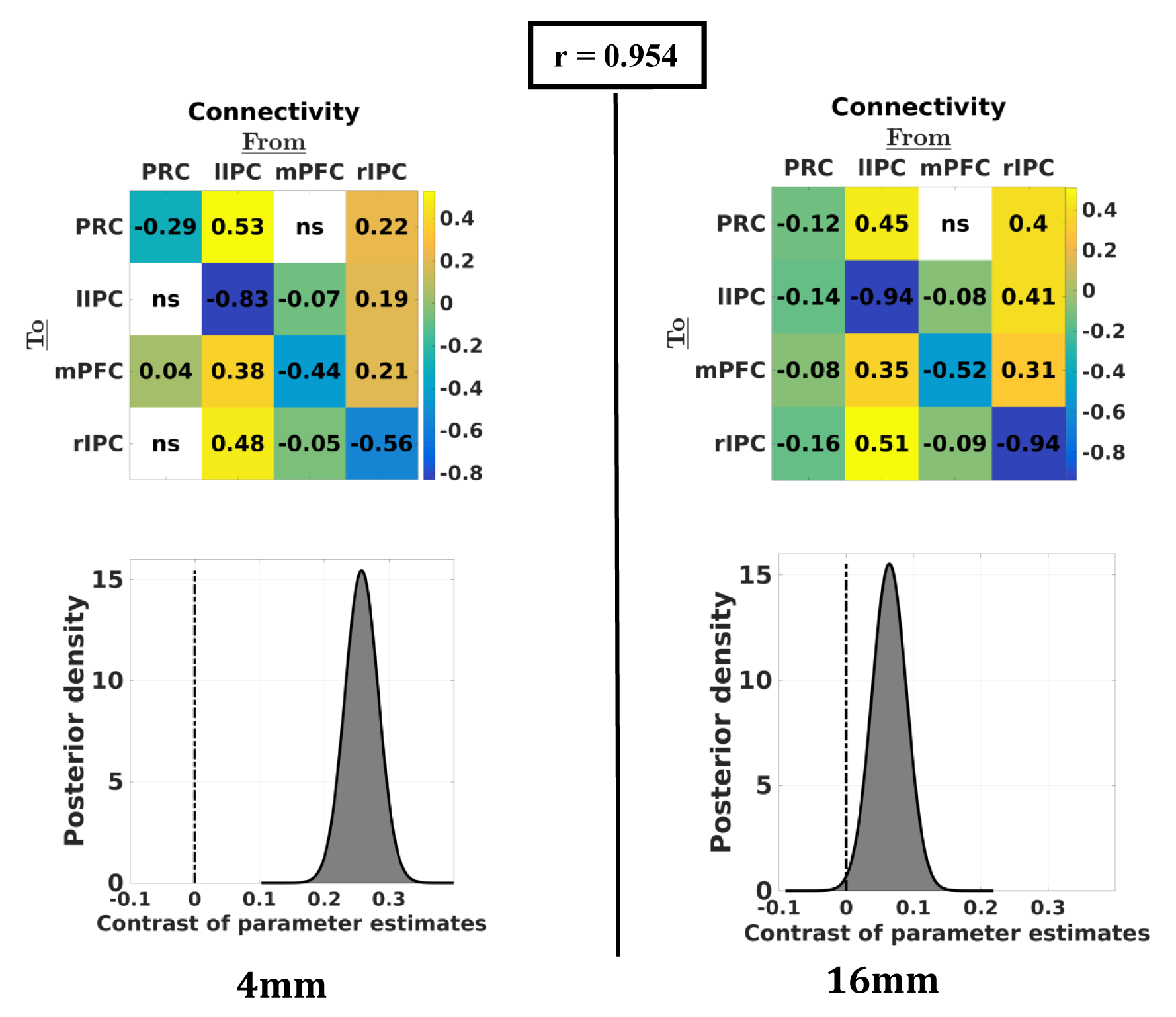
Group-average connectivity for the smallest and largest ROI sizes. Upper panels: effective connectivity matrices for each ROI size at the group-level. Upper number indicates the correlation between (vectorised) matrices. Lower panels: asymmetry of the group-level network (positive = left dominant; negative = right-dominant). Clearly, larger ROI sizes yielded less asymmetry at the group level. However, even at bigger ROIs the network was left-dominant (see posterior probability).

#### Effect of Global Signal Regression

Connectivity with and without GSR is shown in Figure 9. At the group-level, extrinsic connectivity decreased slightly in magnitude after GSR (mean decrease = 0.05Hz), while intrinsic connectivity changed in either a negative or positive direction (depending on the specific region). Importantly, no sign flips were observed. At the subject level, 14 (82%) subjects’ asymmetry patterns were unaffected by GSR, although asymmetry was often less pronounced after GSR. The reliability of connectivity patterns was similar after GSR (average within-subject correlation was 0.39 and 0.37 without and with GSR, respectively).

**Figure 9.**
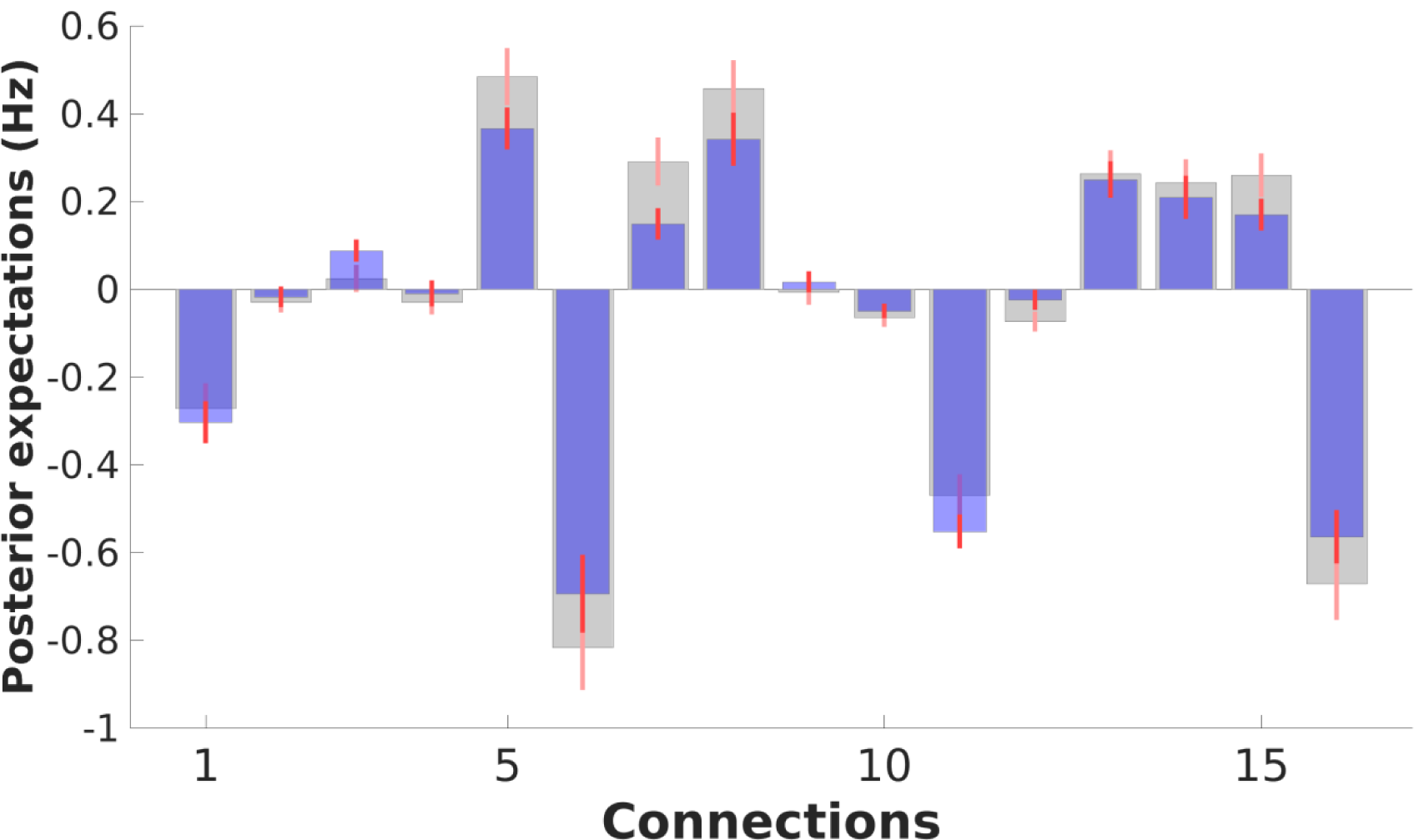
Group-average connectivity with GSR (blue bars), with narrow red bars showing 90% confidence intervals (i.e., Bayesian credible intervals) and without GSR (grey bars). Generally, extrinsic connectivity decreased in magnitude, while intrinsic connections changed in either positive or negative direction. However, no dramatic changes (e.g., significant changes in sign) were found.

## Discussion

In this study, we combined fMRI data from multiple longitudinal studies to investigate within and between subject variability of effective connectivity. Collectively, spectral DCM furnished robust connectivity estimates for each subject, enabled us to track changes in fluctuations across scan sessions, and allowed us to draw conclusions beyond participant groups, scanners, and scanning parameters. Our results revealed that, across datasets, individuals consistently show hemispheric asymmetry of effective connectivity in the default mode network. Previous studies using spectral DCM have also shown some levels of asymmetry at the group-level. Razi et al. (2015) and Zhou et al. (2018) found larger influence from left compared to right parietal cortex, while Sharaev et al. (2016) found the opposite pattern (although the left-right difference was small). Moreover, all studies found that the parietal cortex has a driving influence in the core DMN (Zhou et al., 2018). The present study reproduced the latter result for individual subjects, and suggests that the (small) differences between studies might be attributed to a difference in lateralization patterns of the individual subjects studied. It is worth mentioning that we found that self-inhibition was lowest for the dominant IPC in almost all subjects. This makes sense from a network perspective, since a region that dominates the network should indeed show prolonged (i.e., disinhibited) activity. This observation suggests that the parameters estimated by spectral DCM covary in an intuitive and consistent way.

Lateralization of the default mode network has also been found in other studies using functional connectivity (e.g., Agcaoglu et al., 2015; Nielsen et al., 2013). Agcaoglu et al. (2015) applied independent component analysis to a large group of subjects (n = 603) and found that almost all default mode network components were left lateralized. Similarly, Nielsen et al. (2013) showed that many left-lateralized resting state hubs are part of the default mode network. Our results complement these studies, in showing that hemispheric asymmetry is expressed in terms of effective connectivity, and that the asymmetry is mainly present for interhemispheric connections and connections from lateral to medial areas. Moreover, we have shown that hemispheric asymmetry of effective connectivity is the *main* source of between-subject variability.

The pattern of hemispheric asymmetry was reliable for many subjects. Furthermore, individual connections arising from either right or left IPC showed high sign-stability, which coincided with the individual’s asymmetry. Longitudinal studies assessing the reliability of functional connectivity (e.g., Choe et al., 2015; Gordon et al., 2017) do not often focus on hemispheric asymmetry. However, hemispheric specialization is an important issue in cognitive neuroscience (see, e.g., Hervé et al., 2013). Crucially, a change in DMN lateralization has been associated with psychiatric syndromes (see, e.g., Swanson et al., 2011). The overall stability of the asymmetry of effective connectivity in the DMN could speak to its use as a biomarker for future studies. We also found that some subjects showed more variable hemispheric asymmetry. Within-subject variations in connectivity patterns have also been observed in longitudinal functional connectivity studies. Gordon et al. (2017) found that one specific participant showed considerable lower reliability compared to the others, which they attributed to a higher level of drowsiness. Although this precise subject was left out of the present analyses, it is likely that subject or session specific characteristics (e.g., emotionality) might have caused more variable asymmetry in some subjects compared to others.

To assess the effect of processing methods on connectivity and reliability, the analyses were repeated using global signal regression, varying ROI sizes, and (empirical) Bayesian model reduction. Generally, processing techniques had little effects on our results. Global signal regression had no effect on hemispheric asymmetry for most subjects and did not alter the (within-subject) stability of effective connectivity. Such robustness is quite remarkable, given that the global signal is an important subject of debate in many functional connectivity studies (see e.g., Murphy & Fox, 2017). The robustness of spectral DCM to global confounds is probably explained by the fact that global fluctuations in fMRI signal are modelled explicitly. In other words, parameters representing different sources of noise that are included in DCM can capture fluctuations in global signal that is not mediated by changes in effective connectivity.

Similarly, ROI size had no effect on asymmetry for most subjects and did not have an impact on the reliability of connectivity patterns. Importantly, Bayesian model reduction (BMR; Friston et al., 2016) increased both within- and between-subject consistency of effective connectivity patterns. This suggests that the use of subject and group-specific priors to update the parameters at the session and subject level may enhance reliability, by increasing the probability that parameters are drawn out of local extrema towards the subject or group mean.

Generally, the results raise the question of what might explain the differences between subjects (e.g., hemispheric asymmetry, connection types) and fluctuations within subjects. Variability between subjects might be explained by several factors. First, variability might be related to subject-specific characteristics such as age, gender, and intelligence level. Agcaoglu et al. (2015), for example, showed that age and gender are related to a difference in asymmetry in some resting state networks, most notably the visual network. Similarly, Joliot, Tzourio-Mazoyer, and Mazoyer (2016) showed a relationship between language lateralization and lateralization of resting state connectivity. Our sample comprised participants with a wide age-range (between 24 and 45 years) and was balanced with respect to gender (55% females). Possibly these subjects’ characteristics might have played a role in the extent of asymmetry or at the level of individual connections. Second, differences in scan procedures and sequences might explain observed differences between subjects. Resting state scans are acquired when subjects have either their eyes open or closed; however, no consistent paradigm has been adopted. Studies have shown that a difference in ‘eyes open’ versus ‘eyes closed’ conditions might have an impact on connectivity and reliability during rest (e.g., Zhang et al., 2015; Zou et al., 2015). In our study, the ‘day2day’ dataset was acquired under ‘eyes closed’ conditions and showed a higher proportion of left dominant subjects compared to the ‘MSC’ dataset, which was acquired during ‘eyes open’ conditions. Such procedural differences might thus explain observed differences among subjects. Although several accounts can be offered for between-subject differences, our sample size was too small (n = 17 after exclusion) to support robust explanations. Future studies using datasets with more subjects could try to address possible sources of between-subject variability in effective connectivity, while accounting for within-subject variability.

Fluctuations within individuals might also be explained by several factors. First, fluctuations in effective connectivity might be related to day-to-day changes in mood, behaviour (e.g., amount of sleep), or physiology (e.g., hormonal cycle). Some studies have shown that functional connectivity within the DMN is related to sleepiness (Ward et al., 2013) and is diminished after sleep deprivation (De Havas, Parimal, Soon, & Chee, 2012). Other studies found an influence of female hormones on functional connectivity during rest (e.g., Pletzer, Crone, Kronbichler, & Kerschbaum, 2016). Second, between-subject fluctuations in connectivity might be explained by differences in regional or global noise. Although we did not find a notable difference in parameter estimates with and without GSR, this is not explicit evidence for an influence of global signal on connectivity (or its absence). Similarly, if region-specific noise levels change across sessions, parameter estimates might be affected by conditional dependencies between connectivity and (scale free) noise estimates. Third, Park et al. (2017) have shown that effective connectivity during resting state fluctuates on a short time scale. These faster fluctuations might cause differences among scanning sessions, and therefore explain longitudinal variability in effective connectivity. Indeed, Park et al. (2017) showed that between-session consistency increased when within-session fluctuations were taken into account.

Longitudinal datasets afford the opportunity to test the above hypotheses. Such analyses however fall outside the scope of the present study. Our aim was to provide a framework to test both between- and within-subject variability and consistency of effective connectivity in a single design. The use of PEB, upon which this framework was build, allows researchers to assess relations between connectivity patterns (e.g., asymmetry) and other measures (e.g., physiology), which is of great importance for neuroscience. Future studies could address more refined accounts of between and within subject variability in further detail.

## References

Agcaoglu, O., Miller, R., Mayer, A. R., Hugdahl, K., & Calhoun, V. D. (2015). Lateralization of resting state networks and relationship to age and gender. NeuroImage, 104, 310–325. https://doi.org/10.1016/j.neuroimage.2014.09.001

Buxton, R. B., Wong, E. C., & Frank, L. R. (1998). Dynamics of blood flow and oxygenation changes during brain activation: the balloon model. Magn Reson Med, 39(17), 855–864. https://doi.org/10.1002/mrm.1910390602

Choe, A. S., Jones, C. K., Joel, S. E., Muschelli, J., Belegu, V., Caffo, B. S.,…Pekar, J. J. (2015). Reproducibility and temporal structure in weekly resting-state fMRI over a period of 3.5 years. PLoS ONE, 10(10), 1–29. https://doi.org/10.1371/journal.pone.0140134

Damoiseaux, J. S., Rombouts, S. A. R. B., Barkhof, F., Scheltens, P., Stam, C. J., Smith, S. M., & Beckmann, C. F. (2006). Consistent resting-state networks across healthy subjects. Proceedings of the National Academy of Sciences, 103(37), 13848–13853. https://doi.org/10.1073/pnas.0601417103

De Havas, J. A., Parimal, S., Soon, C. S., & Chee, M. W. L. (2012). Sleep deprivation reduces default mode network connectivity and anti-correlation during rest and task performance. NeuroImage, 59(2), 1745–1751. https://doi.org/10.1016/j.neuroimage.2011.08.026

Filevich, E., Lisofsky, N., Becker, M., Butler, O., Lochstet, M., Martensson, J.,…Kühn, S. (2017). Day2day: Investigating daily variability of magnetic resonance imaging measures over half a year. BMC Neuroscience, 18(1), 1–8. https://doi.org/10.1186/s12868-017-0383-y

Friston, K. J. (2011). Functional and Effective Connectivity: A Review. Brain Connectivity, 1(1), 13–36. https://doi.org/10.1089/brain.2011.0008

Friston, K. J., Harrison, L., & Penny, W. (2003). Dynamic causal modelling. NeuroImage, 19(4), 1273–1302. https://doi.org/10.1016/S1053-8119(03)00202-7

Friston, K. J., Kahan, J., Biswal, B., & Razi, A. (2014). A DCM for resting state fMRI. NeuroImage, 94, 396–407. https://doi.org/10.1016/j.neuroimage.2013.12.009

Friston, K. J., Litvak, V., Oswal, A., Razi, A., Stephan, K. E., Van Wijk, B. C. M.,…Zeidman, P. (2016). Bayesian model reduction and empirical Bayes for group (DCM) studies. NeuroImage, 128, 413–431. https://doi.org/10.1016/j.neuroimage.2015.11.015

Frässle, S., Paulus, F. M., Krach, S., & Jansen, A. (2015). Test-retest reliability of effective connectivity in the face perception network. Human Brain Mapping, 37, 730–744. https://doi.org/10.1002/hbm.23061

Gordon, E. M., Laumann, T. O., Gilmore, A. W., Newbold, D. J., Greene, D. J., Berg, J. J.,…Dosenbach, N. U. F. (2017). Precision Functional Mapping of Individual Human Brains. Neuron, 95(4), 791–807. https://doi.org/10.1016/j.neuron.2017.07.011

Hervé, P.-Y., Zago, L., Petit, L., Mazoyer, B., & Tzourio-Mazoyer, N. (2013). Revisiting human hemispheric specialization with neuroimaging. Trends in Cognitive Sciences, 17(2), 69–80. https://doi.org/10.1016/J.TICS.2012.12.004

Joliot, M., Tzourio-Mazoyer, N., & Mazoyer, B. (2016). Intra-hemispheric intrinsic connectivity asymmetry and its relationships with handedness and language Lateralization. Neuropsychologia, 93, 437–447. https://doi.org/10.1016/j.neuropsychologia.2016.03.013

Laumann, T. O., Gordon, E. M., Adeyemo, B., Snyder, A. Z., Joo, S. J. un, Chen, M. Y.,…Petersen, S. E. (2015). Functional System and Areal Organization of a Highly Sampled Individual Human Brain. Neuron, 87(3), 657–670. https://doi.org/10.1016/j.neuron.2015.06.037

Li, B., Daunizeau, J., Stephan, K. E., Penny, W., Hu, D., & Friston, K. (2011). Generalised filtering and stochastic DCM for fMRI. NeuroImage, 58(2), 442–457. https://doi.org/10.1016/j.neuroimage.2011.01.085

Murphy, K., & Fox, M. D. (2017). Towards a Consensus Regarding Global Signal Regression for Resting State Functional Connectivity MRI. Neuroimage, 154, 169–173. https://doi.org/10.1016/j.neuroimage.2016.11.052

Nielsen, J. A., Zielinski, B. A., Ferguson, M. A., Lainhart, J. E., & Anderson, J. S. (2013). An Evaluation of the Left-Brain vs. Right-Brain Hypothesis with Resting State Functional Connectivity Magnetic Resonance Imaging. PLoS ONE, 8(8). https://doi.org/10.1371/journal.pone.0071275

Park, H.-J., Friston, K. J., Pae, C., Park, B., & Razi, A. (2017). Dynamic effective connectivity in resting state fMRI. NeuroImage. Advance online publication. https://doi.org/10.1016/j.neuroimage.2017.11.033

Pletzer, B., Crone, J. S., Kronbichler, M., & Kerschbaum, H. (2016). Menstrual Cycle and Hormonal Contraceptive-Dependent Changes in Intrinsic Connectivity of Resting-State Brain Networks Correspond to Behavioral Changes Due to Hormonal Status. Brain Connectivity, 6(7), 572–585. https://doi.org/10.1089/brain.2015.0407

Razi, A., & Friston, K. J. (2016). The connected brain: Causality, models, and intrinsic dynamics. IEEE Signal Processing Magazine, 33, 14–35. https://doi.org/10.1109/MSP.2015.2482121

Razi, A., Kahan, J., Rees, G., & Friston, K. J. (2015). Construct validation of a DCM for resting state fMRI. NeuroImage, 106, 1–14. https://doi.org/10.1016/j.neuroimage.2014.11.027

Schuyler, B, Ollinger, J. M., Oakes, T. R., Johnstone, T., & Davidson, R. J. (2010). Dynamic causal modeling applied to fMRI data shows high reliability. NeuroImage, 49(1), 603–611. https://doi.org/10.1016/j.neuroimage.2009.07.015

Sharaev, M. G., Zavyalova, V. V, Ushakov, V. L., Kartashov, S. I., & Velichkovsky, B. M. (2016). Effective Connectivity within the Default Mode Network: Dynamic Causal Modelling of Resting-State fMRI Data. Frontiers in Human Neuroscience, 10 (14). https://doi.org/10.3389/fnhum.2016.00014

Stephan, K. E., Schlagenhauf, F., Huys, Q. J. M., Raman, S., Aponte, E. A., Brodersen, K. H.,…Heinz, (2017). Computational neuroimaging strategies for single patient predictions. NeuroImage, 145, 180–199. https://doi.org/10.1016/j.neuroimage.2016.06.038

Swanson, N., Eichele, T., Pearlson, G., Kiehl, K., Yu, Q., & Calhoun, V. D. (2011). Lateral differences in the default mode network in healthy controls and patients with schizophrenia. Human Brain Mapping, 32(4), 654–64. https://doi.org/10.1002/hbm.21055

Ushakov, V., Sharaev, M. G., Kartashov, S. I., Zavyalova, V. V., Verkhlyutov, V. M., & Velichkovsky, M. (2016). Dynamic Causal Modelling of Hippocampal Links within the Human Default Mode Network: Lateralization and Computational Stability of Effective Connections. Frontiers in Human Neuroscience, 10, 1–14. https://doi.org/10.3389/fnhum.2016.00528

Van Den Heuvel, M. P., Mandl, R. C. W., Kahn, R. S., & Pol, H. E. (2009). Functionally linked resting-state networks reflect the underlying structural connectivity architecture of the human brain. Human Brain Mapping, 30(10), 3127–3141. https://doi.org/10.1002/hbm.20737

Ward, A. M., McLaren, D. G., Schultz, A. P., Chhatwal, J., Boot, B. P., Hedden, T., & Sperling, R. A. (2013). Daytime Sleepiness Is Associated with Decreased Default Mode Network Connectivity in Both Young and Cognitively Intact Elderly Subjects. Sleep, 36(11), 1609– 1615. https://doi.org/10.5665/sleep.3108

Zhang, D., Liang, B., Wu, X., Wang, Z., Xu, P., Chang, S.,…Huang, R. (2015). Directionality of large-scale resting-state brain networks during eyes open and eyes closed conditions. Frontiers in Human Neuroscience, 9 (81), 1–10. https://doi.org/10.3389/fnhum.2015.00081

Zhou, Y., Friston, K. J., Zeidman, P., Chen, J., Li, S., & Razi, A. (2018). The Hierarchical Organization of the Default, Dorsal Attention and Salience Networks in Adolescents and Young Adults. Cerebral Cortex, 28(2), 726–737. https://doi.org/10.1093/cercor/bhx307

Zou, Q., Yuan, B.-K., Gu, H., Liu, D., Wang, D. J. J., Gao, J.-H.,…Zang, Y.-F. (2015). Detecting Static and Dynamic Differences between Eyes-Closed and Eyes-Open Resting States Using ASL and BOLD fMRI. PLOS ONE, 10(3). https://doi.org/10.1371/journal.pone.0121757

